# Targeted profiling of human extrachromosomal DNA by CRISPR-CATCH

**DOI:** 10.1101/2021.11.28.470285

**Authors:** King L. Hung, Jens Luebeck, Siavash R. Dehkordi, Ceyda Coruh, Julie A. Law, William J. Greenleaf, Paul Mischel, Vineet Bafna, Howard Y. Chang

## Abstract

Extrachromosomal DNA (ecDNA) is a common mode of oncogene amplification but is challenging to analyze. Here, we present a method for targeted purification of megabase-sized ecDNA by combining *in-vitro* CRISPR-Cas9 treatment and pulsed field gel electrophoresis of agarose-entrapped genomic DNA (CRISPR-CATCH). We demonstrate strong enrichment of ecDNA molecules containing *EGFR*, *FGFR2* and *MYC* from human cancer cells. Targeted purification of ecDNA versus chromosomal DNA enabled phasing of genetic variants and provided definitive proof of an EGFRvIII mutation on ecDNA and wild-type *EGFR* on chromosomal DNA in a glioblastoma neurosphere model. CRISPR-CATCH followed by nanopore sequencing enabled single-molecule ecDNA methylation profiling and revealed hypomethylation of the *EGFR* promoter on ecDNA compared to the native chromosomal locus in the same cells. Finally, separation of ecDNA species by size and sequencing allowed accurate reconstruction of megabase-sized ecDNA structures with base-pair resolution. CRISPR-CATCH is a new addition to the toolkit for studying focal amplifications in cancer and will accelerate studies aiming to explore the genetic and epigenetic landscapes of ecDNA.

## INTRODUCTION

Oncogene amplification is a key cancer driving mechanism and frequently occurs on circular extrachromosomal DNA (ecDNA). ecDNA oncogene amplifications are present in half of human cancer types and up to one third of tumor samples and are associated with poor patient outcomes^1–3^. Given the prevalence of ecDNA in cancer, there is an urgent need for better characterization of unique genetic and epigenetic features of ecDNA in order to understand how it may differ from chromosomal DNA and obtain clues about how it is formed and maintained in tumors. However, isolation and targeted profiling of megabase-sized, clonal ecDNAs is currently challenging due to their large sizes and sequence complexity, in contrast to small kilobase- and subkilobase-sized DNA circles known as extrachromosomal circular DNA elements (eccDNAs) observed also in non-cancer cells and apoptotic byproducts^4, 5^.

There are currently three main approaches to analyzing sequences of ecDNAs in cancer cells: 1) DNA fluorescence *in situ* hybridization (FISH), 2) bulk whole genome sequencing (WGS), and 3) exonuclease digestion of linear DNA followed by DNA amplification. The first method, DNA FISH, involves arresting cells in metaphase followed by chromosome spreading and hybridization of a DNA probe on a microscope slide. It provides excellent separation of ecDNA and chromosomal DNA signals and has been used to confirm the presence of oncogenes and drug resistance genes on ecDNA. However, this method is low throughput (tens of cells) and provides limited, binary sequence information (a probe either binds or does not bind to DNA). The second method, bulk short- or long-read sequencing, provides much higher sequence resolution. However, sequencing signal represents a combination of all DNA material in a sample, including ecDNA and chromosomal DNA. In addition to the ambiguous origin of sequencing reads, rearranged ecDNA sequences are computationally inferred^1, 6^ but difficult to validate as sequencing reads are far too short to span the entire length of an ecDNA molecule (typically several megabases). Optical mapping (OM) allows analysis of longer DNA molecules (up to several hundred kilobases) by compromising nucleotide-level information but each individual OM molecule is typically shorter than an ecDNA circle^7, 8^. Sequence segments can be computationally “stitched” together to form a list of candidate reconstructed paths, though empirically proving the true ecDNA structure, when possible, is very time-consuming and labor-intensive. The third method, exonuclease treatment combined with DNA amplification, is effective for small DNA circles (up to tens of kilobases; Circle-seq^4, 9^) and was recently applied to ecDNA in cancer cells^10^. It entails magnetic bead-based DNA isolation, treatment with an exonuclease to deplete linear DNA, followed by multiple displacement amplification (MDA). This method requires intact DNA circles and is therefore highly limited by ecDNA size, as megabase-sized DNA molecules are extremely fragile in solution and prone to breakage. Further, this method requires DNA amplification and, therefore, cannot be used for epigenetic analyses. Phi29, the processive MDA polymerase, produces amplicons which are tens of kilobases and thus amplifies small circles via rolling-circle amplification; however, this is currently challenging for megabase-sized ecDNA. Finally, analysis of these enriched ecDNAs by short- or long-read sequencing also suffers from the same read length limitations for amplicon reconstruction.

Here we report a method for purifying megabase-sized ecDNA from cancer cells. DNA amplification is not required; thus, this method allows targeted analyses of both the genetic sequence and epigenetic features of purified ecDNA. We also provide an analytical pipeline for reconstructing amplicon structures *de novo* with high confidence using sequence information of ecDNA species separated by size.

## RESULTS

### Purification and visualization of ecDNA by CRISPR-CATCH

To preserve large intact circular ecDNA, we encapsulated genomic DNA of GBM39 cells (patient-derived glioblastoma neurosphere model containing *EGFR* ecDNA) in agarose plugs (Methods). Fragment size distribution analysis by pulsed field gel electrophoresis (PFGE) showed that virtually all agarose-entrapped genomic DNA containing ecDNA was restricted to either the loading well or the upper compression zone (CZ, region of large DNA molecules; **Extended Data Figure 1a**). ecDNAs are not detectable in the resolution window, suggesting that intact circular ecDNAs do not migrate freely in PFGE as suggested by previous southern blot studies^11–13^. To selectively pull ecDNA into the resolution window of the gel, we pre-incubated GBM39 genomic DNA *in vitro* with CRISPR-Cas9 and a single guide RNA (sgRNA) targeting the *EGFR* locus, an amplified sequence on ecDNA. We reasoned that a single cut would linearize ecDNA, resulting in differential migration in PFGE (**Figure 1a**). We further reasoned that the same single cut in the corresponding chromosomal locus would result in two much larger chromosomal DNA pieces that migrate much more slowly than ecDNA and therefore would not be co-enriched. Cas9 digestion of *EGFR* ecDNA resulted in a prominent band of 1.2-1.37 megabases (**Figure 1b,c**), concordant with the 1.258 Mb amplicon predicted by AmpliconArchitect analysis of bulk WGS and AmpliconReconstructor analysis of OM data^7, 8^. Short-read sequencing of the gel-extracted band confirmed strong enrichment of the expected ecDNA sequence (**Figure 1d,e**), demonstrating that a single cut is sufficient to allow enrichment of ecDNA by PFGE. We refer to this method as CRISPR-CATCH (**C**as9-**A**ssisted **T**argeting of **CH**romosome segments, a term previously coined for a two-cut Cas9 treatment followed by gel extraction for isolating and cloning bacterial chromosomal fragments^14, 15^). CRISPR-CATCH enabled a 30-fold enrichment of the targeted ecDNA (60% of all sequencing reads vs. 2% in WGS), resulting in ultrahigh (∼200x normalized) sequencing coverage (**Figure 1d,e,** ecDNA in **Extended Data Figure 1b**). Simultaneous cleavage of two sgRNA target sites 20 kb away from each other led to loss of the sequence segment between the cut sites, as would be expected given a circular structure and end-to-end junction of the amplified region (**Figure 1f**; ecDNA guides A+B). A single cut in the normal diploid chromosomal *EGFR* locus did not result in a DNA band (as shown in Jurkat cells; **Figure 1c**), further supporting enrichment of ecDNAs in GBM39 cancer cells. To isolate the chromosomal *EGFR* locus, we performed CRISPR-CATCH using two sgRNAs targeting just outside of the amplified region (upstream and downstream; **Figure 1a,b**). This dual-cut strategy resulted in a linear fragment of roughly the same size as the ecDNA molecule and successfully enriched for the chromosomal *EGFR* sequence as demonstrated by increased sequencing coverage around the chromosome-targeting guides (**Figure 1c,f;** chromosomal DNA, **Extended Data Figure 1b**). Chromosomal gel bands appeared much fainter than ecDNA bands (**Figure 1c**), consistent with the fact that ecDNAs exist in higher copy numbers than the chromosomal locus in GBM39 cells. Sequencing coverage analysis further validated enrichment of ecDNA versus chromosomal DNA alleles (**Extended Data Figure 1c,d**). Together, these results showed that CRISPR-CATCH can be used to isolate megabase-sized ecDNA molecules and corresponding chromosomal locus from the same cancer cell sample. CRISPR-CATCH also provides the first empirical proof for amplicon size predictions made by bulk sequencing analysis via migration patterns in PFGE.

**Figure 1.**
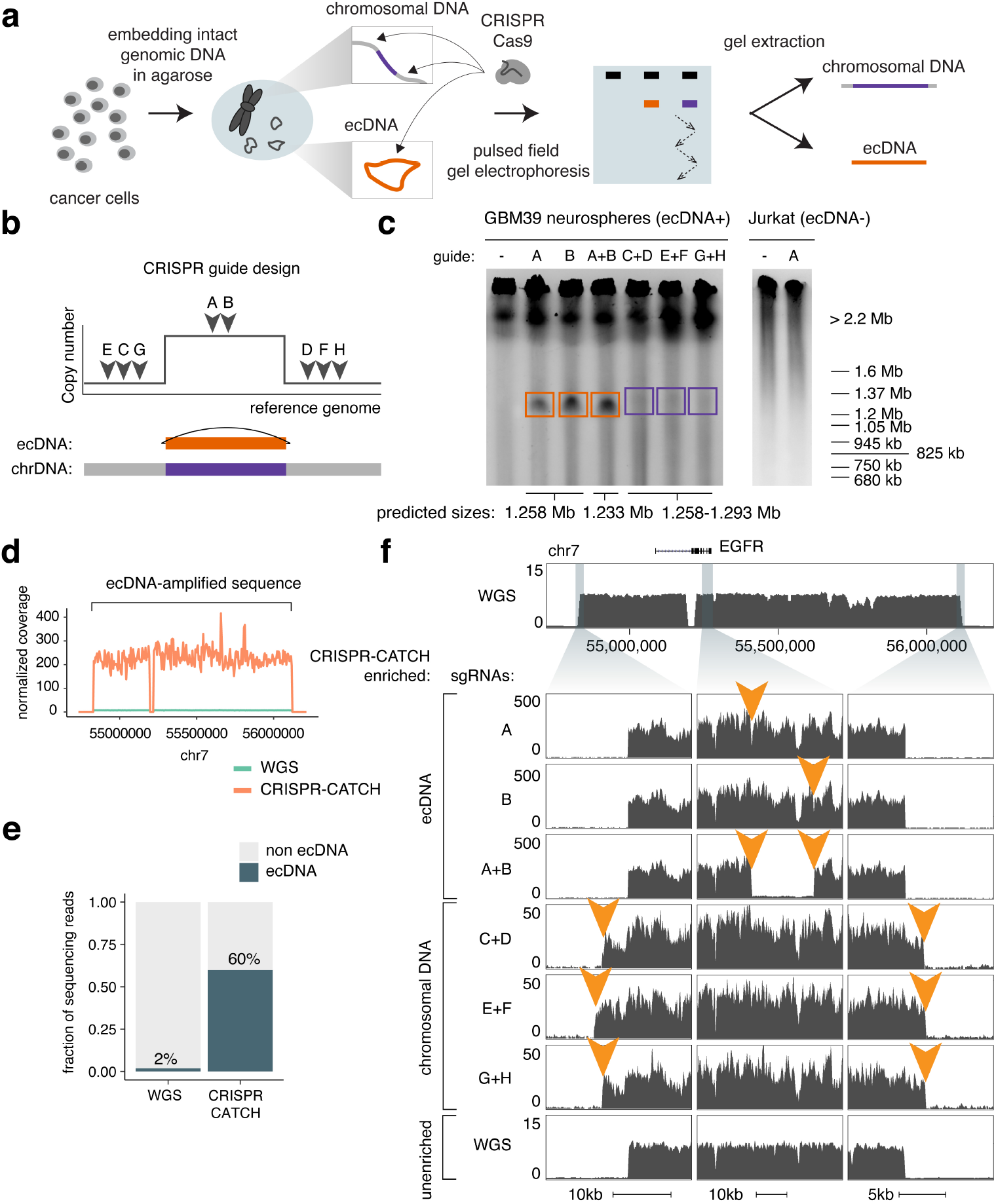
Isolation of megabase-sized ecDNA and its native chromosomal locus from the same cancer cell sample by CRISPR-CATCH. **(a)** Experimental workflow for purification of ecDNA and its corresponding chromosomal locus from the same cell sample. **(b)** Design of CRISPR sgRNAs for linearizing ecDNA circles or extracting the native chromosomal locus. **(c)** Representative PFGE images showing linearized ecDNA molecules and the chromosomal locus after treatment with indicated guides (Methods, raw gel images in **Supplementary** Figure 1, guide sequences in **Supplementary Table 1**). Boxed regions indicate parts of the gel that were extracted for DNA isolation. **(d)** Normalized short-read sequencing coverage of the expected ecDNA locus in unenriched WGS or after CRISPR-CATCH (guide A). **(e)** Fraction of total sequencing reads aligning to the expected ecDNA locus in unenriched WGS or after CRISPR-CATCH (guide A). **(f)** Sequencing tracks showing coverages for purified ecDNA and its chromosomal locus at the zoomed-in locations compared to WGS. Orange arrows indicate locations of sgRNA targets.

### Phasing oncogenic mutations and single-nucleotide variants on ecDNA

Next, we performed targeted analysis of the genetic sequences of ecDNA and chromosomal DNA containing the same oncogene locus (**Figure 2a**). Using ecDNA and chromosomal DNA molecules containing the *EGFR* locus isolated using CRISPR-CATCH, we first identified structural variants using short-read sequencing data. GBM39 cells were previously shown to harbor the EGFRvIII deletion, an activating *EGFR* mutation^7, 8, 16^. Importantly, sequencing coverage combined with breakpoint analysis of CRISPR-CATCH data revealed that the EGFRvIII mutation is predominantly found on ecDNA, while the chromosomal locus mainly contains full-length *EGFR* (**Figure 2b**). Full-length EGFR appeared at ∼75% in the chromosomal fraction, consistent with the level of chromosomal DNA enrichment (**Figure 2b**, **Extended Data Figure 1d**). This observation suggests that the EGFRvIII mutation occurred after ecDNA formation and was strongly selected subsequently (**Figure 2c**). This finding supports previous studies suggesting that ecDNA may help cancer cells adapt to selective pressure and harbor unique genetic alterations^6, 17, 18^.

**Figure 2.**
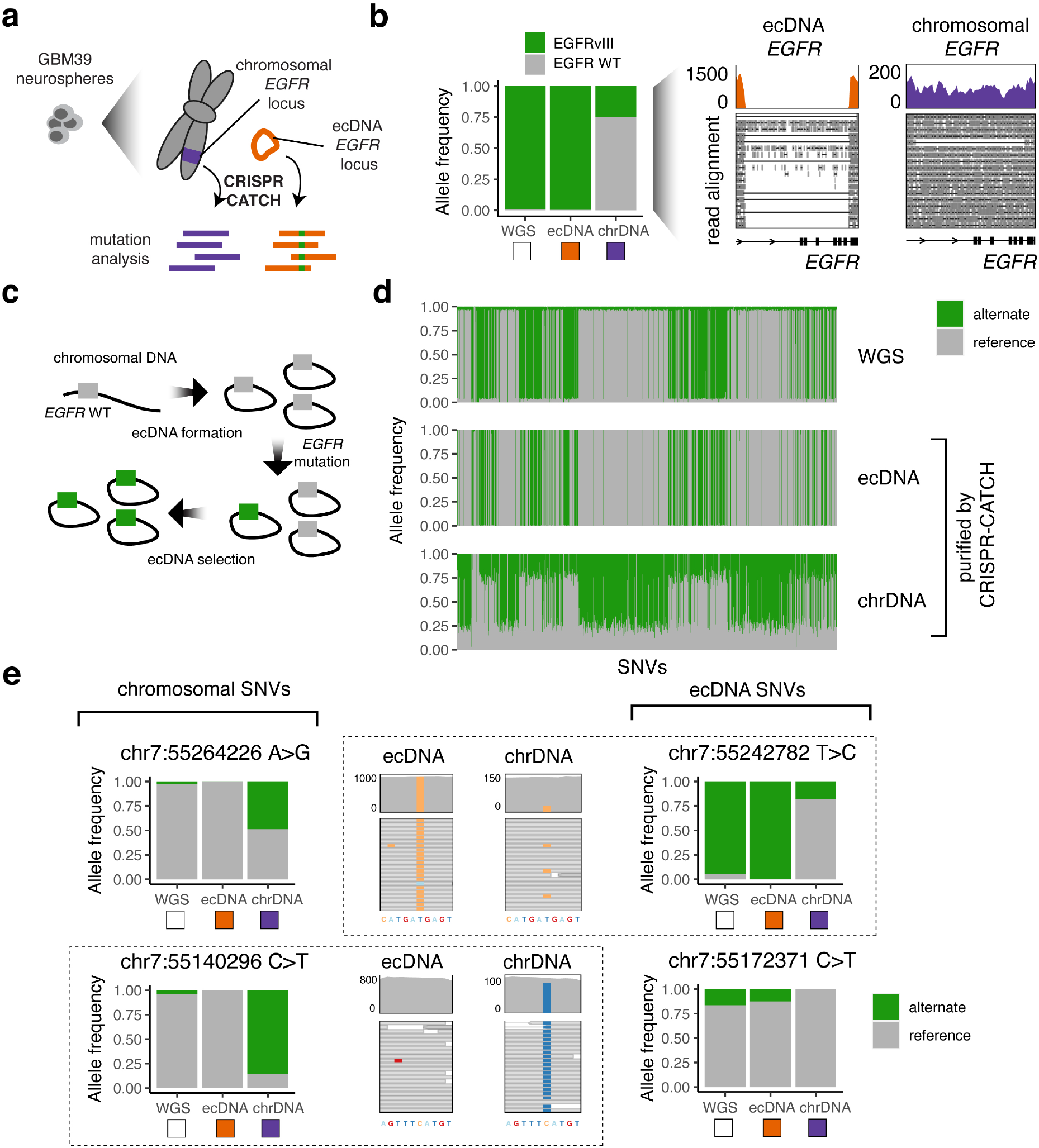
Phasing of SNVs and SVs for ecDNA and its native chromosomal locus. **(a)** Isolation of ecDNA (guide A) and the corresponding chromosomal locus (guides E+F) from GBM39 neurospheres by CRISPR-CATCH followed by mutation analysis using short-read sequencing. **(b)** Left: allele frequencies of the EGFRvIII mutant on ecDNA and chromosomal DNA. Right: sequencing coverage and junction reads supporting the EGFRvIII mutation and wild type. **(c)** EGFRvIII mutation likely occurred after ecDNA formation and became the predominant version of *EGFR* on ecDNA molecules as a result of selection. **(d)** Allele frequencies of SNVs identified in the ecDNA-amplified region and its native chromosomal locus. **(e)** Left and right: examples of ecDNA and chromosomal SNVs with various allele frequencies. Middle: sequencing reads supporting SNV identification. Each dashed box groups data for the same SNV.

We then assessed the frequencies of single-nucleotide variants (SNVs) found on purified ecDNA and chromosomal DNA. We identified SNVs most frequently observed on ecDNAs, including clonal and subclonal variants (**Figure 2d,e**). Notably, we also found a number of unique high-allele-frequency SNVs located in the chromosomal *EGFR* locus, suggesting that the ecDNA and chromosomal loci may have diverged relatively early in the mutational timeline (**Figure 2d,e**). Similar to the EGFRvIII analysis, unique SNVs located in the chromosomal fraction exhibited allele frequencies of 70-75%, consistent with the level of chromosomal DNA enrichment (**Figure 2d**, **Extended Data Figure 1d**). Low-frequency, subclonal ecDNA and chromosomal DNA mutations are indistinguishable in bulk WGS data but can be clearly phased by CRISPR-CATCH (**Figure 2e**).

### Single-molecule DNA methylation profile of purified ecDNA

We then examined the feasibility of analyzing epigenomic profiles of ecDNA using CRISPR-CATCH. After ecDNA purification as before, we performed nanopore sequencing to obtain single-molecule sequence information and DNA cytosine methylation (5mC) profiles. We analyzed 5mC-CpG methylation of purified ecDNA as a proof of concept, and observed a strong anti-correlation of 5mC with chromatin accessibility based on bulk ATAC-seq, validating the identification of regulatory elements (Methods, **Figure 3a,b, Extended Data Figure 2a**). We also purified the corresponding *EGFR* chromosomal locus in GBM39 cells and analyzed its DNA methylation profile (**Figure 3a,b**). We observed reduced DNA methylation at regulatory elements on ecDNA compared to the same elements on chromosomal DNA, suggesting altered gene regulation (top 50 ATAC-seq peaks; **Figure 3c**). The four regions that lost 5mC on ecDNA compared to its chromosomal locus in the same cells were all gene promoters, including that of the *EGFR* oncogene (Methods, **Figure 3d,e, Extended Data Figure 2b**). The pattern of hypomethylation corresponded to nucleosome positions shown by MNase-seq, implying a more active chromatin state on ecDNA (**Figure 3e, Extended Data Figure 2c**)^19, 20^. Finally, single-molecule analysis of purified ecDNA at the *EGFR* promoter showed hypomethylation at the *EGFR* promoter and co-occurrence of methylation spanning hundreds of CpG sites around the region on the same molecules (285 CpG sites, **Figure 3f**). Together, these data show that gene promoters on ecDNA may have increased activities compared to the corresponding chromosomal locus on a single-molecule level and demonstrate that CRISPR-CATCH can be used to measure epigenomic features of ecDNA.

**Figure 3.**
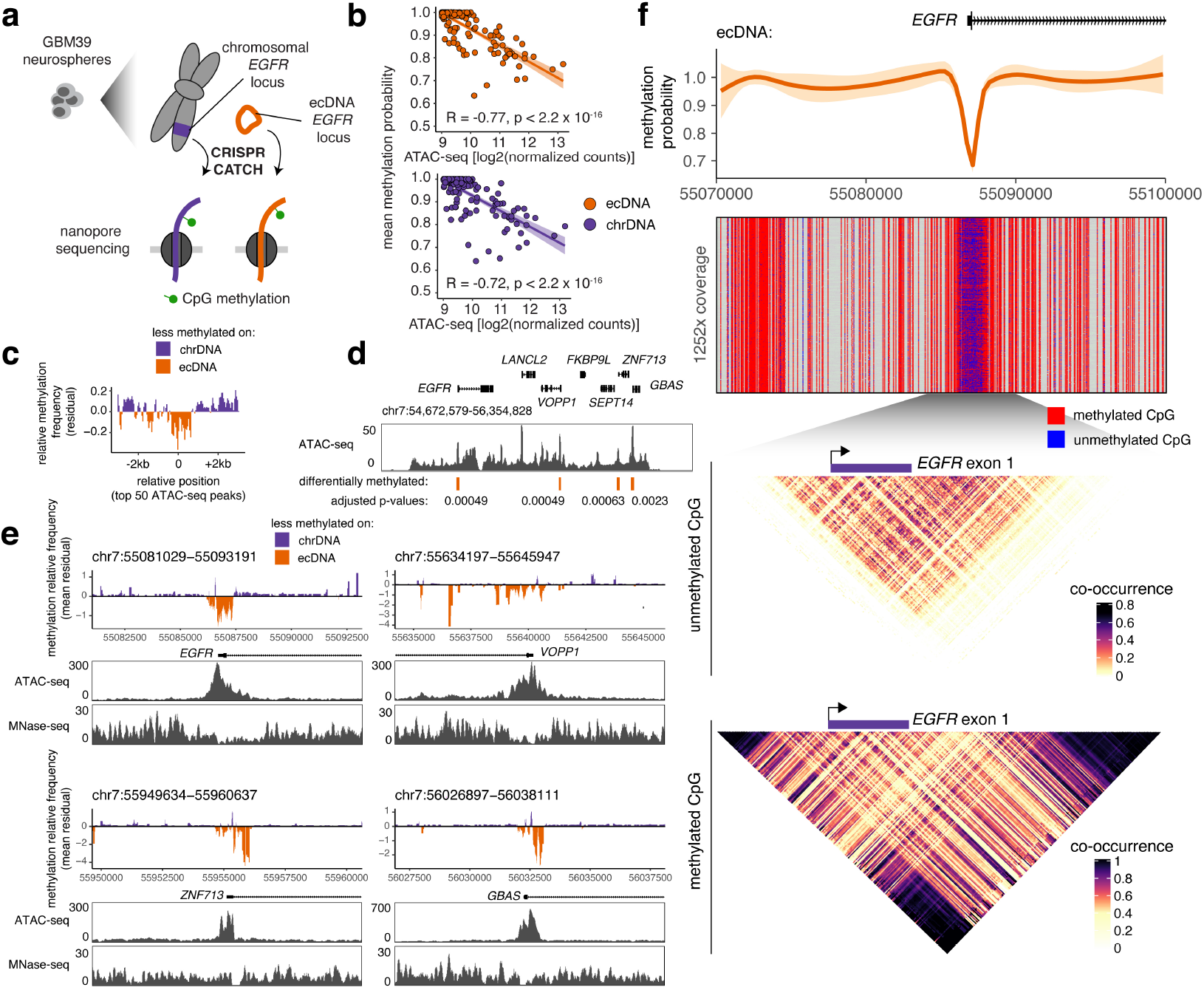
Comparison of CpG methylation statuses of ecDNA and its native chromosomal locus in the same cells. **(a)** Isolation of ecDNA (guide A) and the corresponding chromosomal locus (guides E+F) from GBM39 neurospheres by CRISPR- CATCH followed by detection of 5mC-CpG methylation by nanopore sequencing. **(b)** Negative correlation between mean methylation probabilities of ATAC-seq peaks and their ATAC-seq signals (Pearson’s R, two-sided test). **(c)** Aggregated levels of relative CpG methylation of ecDNA compared to the chromosomal locus at top 50 ATAC-seq peaks in the ecDNA-amplified region. Mean methylation frequencies were calculated in 100-bp windows sliding every 10 bp. Relative frequencies were quantified from standardized residuals for a linear regression model for mean frequencies on ecDNA vs chromosomal DNA (Methods). **(d)** Bulk ATAC-seq track with differentially methylated regions annotated (Methods; p values were Benjamini-Hochberg adjusted; regions with p < 0.005 were considered significant). **(e)** Relative CpG methylation of ecDNA compared to the chromosomal locus in differential regions and concordance with accessibility by ATAC-seq and nucleosome positioning by MNase-seq. Mean methylation frequencies were calculated in 100-bp windows sliding every 10 bp. Relative frequencies were quantified from standardized residuals for a linear regression model for mean frequencies on ecDNA vs chromosomal DNA (Methods). **(f)** From top to bottom: Loess-smoothed methylation probability around the *EGFR* promoter; nanopore sequencing reads showing CpG methylation calls (grey denotes regions with no CpG sites); heatmap showing co- occurrence probabilities of unmethylated CpG sites on the same molecules; heatmap showing co-occurrence probabilities of methylated CpG sites on the same molecules (Methods). Raw PFGE image of purified DNA products is in **Supplementary** Figure 1.

### *De novo* reconstruction of ecDNA amplicon structures

Many cancer cells contain ecDNAs with more complex, heterogeneous structures, including multiple sequence rearrangements and more than one circle species^6^. We reasoned that CRISPR-CATCH may provide direct evidence of molecule size and amplicon-phased structural information for these complex amplicons, and that this information can be used to computationally reconstruct ecDNA with higher confidence. To this end, we developed an analytical pipeline for *de novo* amplicon reconstruction from CRISPR-CATCH data (Methods, **Figure 4a**). We modified and adopted AmpliconArchitect^6^ for generating a copy-number-aware breakpoint graph for each isolated amplicon. Next, we implemented a new method for extracting ecDNA candidate paths from the graph, called **C**andidate **AM**plicon **P**ath **E**numerato**R** (CAMPER). Candidate ecDNA structures were generated from the breakpoint graph, estimated multiplicity of genomic segments and molecular size based on PFGE using a depth-first search (DFS) approach (Methods). Finally, quality estimates of resulting structures were produced for filtering out any low-confidence reconstructions in the case of low-quality gel extractions (e.g., incompletely separated ecDNA species) or undetectable breakpoints from sequencing, etc. As validation, we reconstructed the 1.258 Mb circular ecDNA circle encoding *EGFR* in GBM39 cells using this workflow, yielding a structure fully consistent with previous reports using WGS and optical mapping^7, 8^ (**Extended Data Figure 3**). To further demonstrate the utility of this tool, we applied this pipeline to a stomach cancer cell line, SNU16, which contains multiple ecDNA species with *MYC*, *FGFR2* and additional sequences connected by complex structural rearrangements (**Extended Data Figure 4a**)^21^. CRISPR-CATCH using guides targeting the *MYC* or *FGFR2* amplicon resulted in multiple visible bands in PFGE (**Figure 4b**), revealing extensive molecular heterogeneity of ecDNAs. Gel-extracted ecDNAs were multiplexed for sequencing. In 4 of 23 libraries, short-read sequencing of the CRISPR-CATCH-purified band was sufficient to enable end-to-end, megabase-scale reconstruction of the ecDNA sequence. 5 libraries corresponded to the compression zone and showed very low levels of ecDNA enrichment, suggesting that the true ecDNA sizes are smaller than 2.2 Mb (bands a,e,h,o,r, **Figure 4b**, **Extended Data Figure 5**). In the remaining cases, large amplicon sequences were enriched, but one or more missing edges prevented unambiguous amplicon resolution (**Figure 4b**, **Extended Data Figure 5**). From these data, we reconstructed three unique ecDNAs containing *MYC* or *FGFR2*: a 1.604 Mb *FGFR2* ecDNA that was reconstructed from two independent CRISPR-CATCH treatments (using sgRNAs with cut sites > 300 kb apart), a smaller *FGFR2* ecDNA species that was 228 kb, and a 622 kb *MYC* ecDNA containing sequences originating from chromosomes 8 and 11 (**Figure 4c-e**). All reconstructions from CRISPR-CATCH data passing quality filters were supported by contigs assembled from optical mapping data (N50 50 Mb) provided to AmpliconReconstructor^7^, further validating their structures (Methods, **Figure 4c-e**).

**Figure 4.**
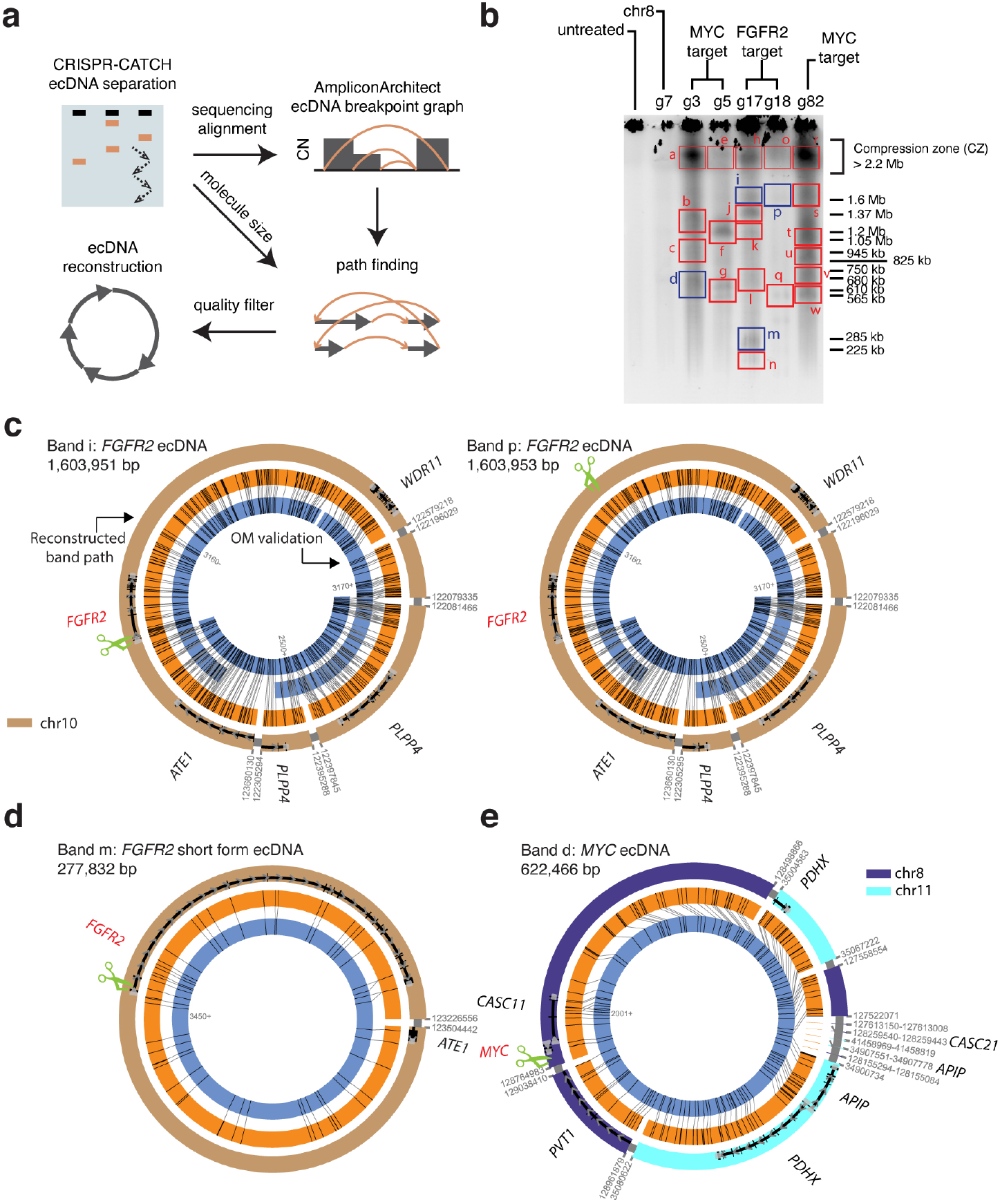
Reconstruction of multiple ecDNA species in the stomach cancer cell line SNU16. **(a)** Analysis of ecDNA structure using CRISPR-CATCH. ecDNA species are separated by size in PFGE and sequenced. AmpliconArchitect generates CN-aware breakpoint graphs, which are used in combination with molecule sizes from PFGE to find paths and identify candidate ecDNA structures. (**b)** PFGE image for SNU16 after treatment with independent sgRNAs targeting either the *FGFR2* or *MYC* locus (guide sequences in **Supplementary Table 1,** raw gel image in **Supplementary** Figure 1). Bands passing all quality filters are shown in blue. **(c-e)** ecDNA reconstructions using CRISPR-CATCH data (outer rings; thin grey bands mark connections between sequence segments). Optical mapping patterns (orange rings) and assembled contigs (blue rings, contig IDs indicated) validated CRISPR-CATCH reconstructions. Green scissors mark sgRNA target sites. Two equivalent *FGFR2* ecDNA structures were reconstructed from bands “i” and “p” from independent sgRNA treatments as shown in panel **(c)**. A short- form *FGFR2* ecDNA reconstructed from band “m” is shown in panel **(d)**. A *MYC* ecDNA reconstructed from band “d” containing sequences from chromosomes 8 and 11 is shown in panel **(e)**.

Breakpoint graphs of ecDNA species were greatly simplified by CRISPR-CATCH because each amplicon could be separately reconstructed and was not intermixed with all other amplicons (**Extended Data Figure 4b**). Together, these data demonstrate the utility of CRISPR-CATCH as a method for disambiguating ecDNA structure and size, particularly when a heterogeneous mixture of ecDNAs is present. The method also aids in accurate amplicon reconstruction orthogonal to contig assembly from bulk DNA.

## DISCUSSION

By exploiting the distinctive PFGE migration pattern of large circular ecDNA, we show that ecDNA can be purified from human cancer cells and separated by size using CRISPR-CATCH. This method enables targeted analyses of ecDNA sequences and epigenomic features that could not be previously achieved. CRISPR-CATCH also makes it possible to directly compare ecDNA and the corresponding chromosomal locus in the same cell sample by physically separating them. It is now possible to obtain allele-specific information of ecDNA versus chromosomal DNA without relying on SNVs. Further, the ability to phase SNVs by CRISPR-CATCH also enables identification of sequencing signal originating from ecDNA in order to obtain allele-specific information (e.g. in bulk RNA-seq data).

The scope and challenge of ecDNA isoforms were not fully appreciated in the past. With the ability to separate ecDNA from the rest of the genome and accurately reconstruct amplicon structures, CRISPR-CATCH may be applied to future studies on cancer cells during early formation of ecDNA, cells evolving under chemotherapeutic or other selective pressures, and in other settings where changes in genetic and chromatin features of ecDNA are hypothesized to contribute to cancer cell evolution. As ecDNA often exhibits tremendous structural heterogeneity, CRISPR-CATCH opens up a new window into deciphering intratumoral genetic heterogeneity in cancer. The ability to separate ecDNAs by size may provide increased structural resolution to other types of analysis, such as single-cell sequencing, in which heterogeneous mixes of ecDNA structures are computationally inferred but difficult to resolve confidently. These future applications of CRISPR-CATCH may also address how ecDNA and chromosomal DNA diverge as they evolve separately and under different kinetics. We note that tandem duplications on chromosomal DNA (e.g. homogeneously staining regions) can also be isolated by CRISPR-CATCH with a single guide. Thus, CRISPR-CATCH should be used to complement additional methods like metaphase FISH to verify the source of isolated DNA.

We demonstrate that CpG methylation can be measured from purified ecDNA molecules. Past studies have shown that cells containing ecDNA express amplified genes at higher levels than cells containing linear amplifications, and that the ecDNA oncogene locus is more accessible than other loci on linear DNA by bulk ATAC-seq^1, 8^. Our comparison of ecDNA versus chromosomal DNA encoding the same gene loci from the same cells showed that gene promoters on circular ecDNA are less methylated than the same promoters on linear chromosomal DNA, suggesting that ecDNA enables more active transcription. In principle, CRISPR-CATCH may be coupled to several genomic assays to understand key chromatin-templated processes on ecDNA such as transcription, DNA replication, and repair^22–24^. CRISPR-CATCH presents an opportunity for a multitude of molecular studies which will help elucidate how ecDNA oncogene amplifications are regulated in cancer cells.

## METHODS

### Cell Culture

GBM39 neurospheres were derived from patient tissue as previously described^8^. All other cell lines used were obtained from ATCC. GBM39 cells were maintained in DMEM/Nutrient Mixture F-12 (DMEM/F12 1:1; Gibco, Cat# 11320-082), B-27 Supplement (Gibco, Cat# 17504044), 1% penicillin-streptomycin (pen-strep; Thermo Fisher, Cat# 15140-122), human epidermal growth factor (EGF, 20 ng/ml; Sigma-Aldrich, E9644), human fibroblast growth factor (FGF, 20 ng/ml; Peprotech) and Heparin (5 ug/ml; Sigma-Aldrich, Cat# H3149-500KU). SNU16 cells were maintained in DMEM/F12 supplemented with 10% FBS and 1% pen-strep. All cells were cultured at 37°C with 5% CO2. All cell lines tested negative for mycoplasma contamination.

### Whole-Genome Sequencing

Whole genome sequencing (WGS) data from bulk GBM39 cells were generated by a previously published study^8^ and raw fastq reads obtained from the NCBI Sequence Read Archive, under BioProject accession PRJNA506071. Reads were trimmed of adapter content with Trimmomatic^25^ (version 0.39), aligned to the hg19 genome using BWA MEM^26^ (0.7.17-r1188), and PCR duplicates removed using Picard’s MarkDuplicates (version 2.25.3). WGS data from bulk SNU16 cells were previously generated (SRR530826, Genome Research Foundation).

### ecDNA isolation by CRISPR-CATCH

Genomic DNA was embedded in agarose plugs using a modified protocol based on guidelines from the manufacturer of the CHEF Mapper XA System (Bio-Rad Laboratories) as previously described^27^. Briefly, molten 1% certified low melt agarose (Bio-Rad, 1613112) in PBS was equilibrated to 45°C. 1 million cells were pelleted per condition, washed twice with cold 1X PBS, resuspended in 30 ul PBS, and briefly heated to 37°C. 30 ul agarose solution was added to cells, mixed, transferred to a plug mold (Bio-Rad Laboratories, Cat #1703713) and incubated on ice for 10 minutes. Solid agarose plugs containing cells were ejected into 1.5 ml Eppendorf tubes, suspended in buffer SDE (1% SDS, 25mM EDTA at pH 8.0) and placed on shaker for 10 minutes. The buffer was removed and buffer ES (1% N-laurolsarcosine sodium salt solution, 25 mM EDTA at pH 8.0, 50ug/ml proteinase K) was added. Agarose plugs were incubated in buffer ES at 50°C overnight. On the following day, proteinase K was inactivated with 25 mM EDTA with 1 mM PMSF for 1 hour at room temperature with shaking. Plugs were then treated with RNase A (1mg/ml) in 25 mM EDTA for 30 minutes at 37°C, and washed with 25 mM EDTA with a 5-minute incubation. Plugs not directly used for ecDNA purification were stored in 25 mM EDTA at 4°C.

To perform *in-vitro* Cas9 digestion, agarose plugs containing DNA were washed three times with 1X NEBuffer 3.1 (New England BioLabs) with 5-minute incubations. Next, DNA was digested in a reaction with 30 nM single-guide RNA (sgRNA, Synthego) and 30 nM spCas9 (New England BioLabs, M0386S) after pre-incubation of the reaction mix at room temperature for 10 minutes. To make two cuts on the native chromosomal locus, 15 nM of each sgRNA was added to the reaction. Cas9 digestion was performed at 37°C for 4 hours, followed by overnight digestion with 3 ul proteinase K (20mg/ml) in a 200 ul reaction. On the following day, proteinase K was inactivated with 1 mM PMSF for 1 hour with shaking. plugs were then washed with 0.5X TAE buffer three times with 5-minute incubations. Plugs were loaded into a 1% certified low melt agarose gel (Bio-Rad, 1613112) in 0.5X TAE buffer with ladders (CHEF DNA Size Marker, 0.2–2.2 Mb, S. cerevisiae Ladder: Bio-Rad, 1703605; CHEF DNA Size Marker, 1–3.1 Mb, H. wingei Ladder: Bio-Rad, 1703667) and PFGE was performed using the CHEF Mapper XA System (Bio-Rad) according to the manufacturer’s instructions and using the following settings: 0.5X TAE running buffer, 14°C, two-state mode, run time duration of 16 hours 39 minutes, initial switch time of 20.16 seconds, final switch time of 2 minutes 55.12 seconds, gradient of 6 V/cm, included angle of 120°, and linear ramping. Gel was stained with 3X Gelred (Biotium) with 0.1 M NaCl on a rocker for 30 minutes covered from light and imaged. Bands were then extracted and DNA was purified from agarose blocks using beta-Agarase I (New England BioLabs, M0392L) following the manufacturer’s instructions.

### Short-read sequencing of DNA isolated by CRISPR-CATCH

To perform short-read sequencing on DNA isolated by CRISPR-CATCH, we first transposed it with Tn5 transposase produced as previously described^28^, in a 50 ul reaction with TD buffer^29^, 10 ng DNA and 1 ul transposase. The reaction was performed at 37°C for 5 minutes, and transposed DNA was purified using MinElute PCR Purification Kit (Qiagen, 28006). Libraries were generated by 7 rounds of PCR amplification using NEBNext High-Fidelity 2X PCR Master Mix (NEB, M0541L), purified using SPRIselect reagent kit (Beckman Coulter, B23317) with double size selection (0.8X right, 1.2X left) and sequenced on the Illumina Miseq platform or on an Illumina NovaSeq 6000. For GBM39 enrichment and mutation analyses in Figures 1 and 2, a 1.2X left-side selection was performed using SPRIselect. Sequencing data were processed as described above for WGS.

### Genetic Variant Analyses

SNVs were identified using GATK (version 4.2.0.0)^30^ from short-read sequencing data as follows. First, base quality score recalibration was performed on bam files (generated as described above) using gatk BaseRecalibrator followed by gatk ApplyBQSR. Covariates were analyzed using gatk AnalyzeCovariates. SNVs were called using gatk Mutect2 from the recalibrated bam files, and SNVs were filtered using gatk FilterMutectCalls. Finally, vcf files were converted to table format using gatk VariantsToTable with the following parameters: “-F CHROM -F POS -F REF -F ALT -F QUAL -F TYPE -GF AD -GF GQ -GF PL -GF GT”. Mutation variant allele frequencies (VAFs) were calculated by dividing alternate allele occurrences by the sum of reference and alternate allele occurrences. SNVs which had coverage depth of 5 or less, had a VAF of 1, or were not detected in WGS were filtered out. Read alignment was visualized using Gviz in R.

SVs from short-read sequencing were identified with DELLY^31^ (version 0.8.7; using Boost version 1.74.0 and HTSlib version 1.12) using the delly call command. BCF files were converted to VCF using bcftools view in Samtools^32^. VAFs were calculated using both imprecise and precise variants. Read alignment was visualized using Gviz in R.

### Nanopore Sequencing and 5mC methylation calling

DNA isolated by CRISPR-CATCH was directly used without amplification for nanopore sequencing. Sequencing libraries were prepared using the Rapid Sequencing Kit (Oxford Nanopore Technologies, SQK-RAD004) according to the manufacturer’s instructions. Sequencing was performed on a MinION (Oxford Nanopore Technologies).

Bases were called from fast5 files using guppy (Oxford Nanopore Technologies, version 5.0.16) within Megalodon (version 2.3.3) and DNA methylation status was determined using Rerio basecalling models with the configuration file “res_dna_r941_min_modbases-all-context_v001.cfg” and the following parameters: “-- outputs basecalls mod_basecalls mappings mod_mappings mods per_read_mods -- mod-motif Z CG 0 --write-mods-text --mod-output-formats bedmethyl wiggle --mod-map- emulate-bisulfite --mod-map-base-conv C T --mod-map-base-conv Z C”. Methylation calls on single molecules were visualized using Integrative Genome Viewer (IGV, version 2.11.1) in bisulfite mode.

To quantify 5mC-CpG methylation levels across an entire locus, rolling averages of CpG methylation percentages were calculated using a window of 100 bp sliding every 10 bp (unless otherwise specified). Rolling averages of ecDNA and the native chromosomal locus were linearly regressed using the lm function in R. Standardized residual for the linear regression for each window was calculated using the rstandard function to represent relative methylation frequencies on ecDNA compared to chromosomal DNA. To identify accessible regions which are differentially methylated on ecDNA, we first filtered on ATAC-seq peaks which had log-normalized coverage above 9 (calculated by DESeq2 as described in the ATAC-seq section below; normalized coverage for each peak was divided by peak width after adding 1, scaled to 500 and log2-transformed). Next, methylation sites with coverage above 5 for both the purified ecDNA and chromosomal locus and overlapping filtered ATAC-seq peaks were linearly regressed using the lm function in R. Standardized residual for the linear regression for each CpG site was calculated using the rstandard function. For each ATAC-seq peak, a z score was calculated using the formula z = (x-m)/S.E., where x is the mean CpG residual within the peak, m is the mean residual of all CpG sites, and S.E. is the standard error calculated from the standard deviation of all CpG sites divided by the square root of the number of CpG sites within the peak. Z scores were used to compute two-sided p values using the normal distribution function, which were adjusted with p.adjust in R (version 3.6.1) using the Benjamini-Hochberg Procedure.

To quantify co-occurrence of methylated or unmethylated CpGs on single molecules, methylation calls on the “+” strand were offset by 1 bp to match the locations of the corresponding CpG sites on the “-” strand. CpG sites where the base probabilities of methylation were above 0.7 were categorized as methylated, and sites where the base probabilities of unmodified CpG were above 0.7 were categorized as unmethylated. For each pair of CpG sites, co-occurrence was calculated by number of co-occurrences of methylated or unmethylated CpGs on the same nanopore sequencing reads divided by total number of occurrences in which the two CpG sites can be successfully categorized as either methylated or unmethylated.

### ATAC-seq

ATAC-seq data for GBM39 were generated by a previously published study^8^ and raw fastq reads obtained from the NCBI Sequence Read Archive, under BioProject accession PRJNA506071. Adapter-trimmed reads were aligned to the hg19 genome using Bowtie2 (2.1.0). Aligned reads were filtered for quality using samtools (version 1.9)^32^, duplicate fragments were removed using Picard’s MarkDuplicates (version 2.25.3), and peaks were called using MACS2 (version 2.2.7.1)^33^ with a q-value cut-off of 0.01 and with a no-shift model. Peaks from replicates were merged, read counts were obtained using bedtools (version 2.30.0)^34^ and normalized using DESeq2 (using the “counts” function in DESeq2 with normalized = TRUE; version 1.26.0)^35^.

### MNase-seq

MNase-seq data for GBM39 were generated by a previously published study^8^ and raw fastq reads obtained from the NCBI Sequence Read Archive, under BioProject accession PRJNA506071. Reads were trimmed of adapter content with Trimmomatic^25^ (version 0.39), aligned to the hg19 genome using BWA MEM^26^ (0.7.17-r1188), and PCR duplicates removed using Picard’s MarkDuplicates (version 2.25.3). Coverage of nucleosome midpoints was obtained using bamCoverage from deepTools (version 3.5.1) with the following parameters: “--MNase --binSize 1”.

### *De Novo* Amplicon Reconstruction

Using short-read sequencing data from CRISPR-CATCH with double size selection as described above, we implemented new strategies and modified existing methods^6^ to resolve ecDNA structures. Broadly, the methods involved seven steps. The last six steps are available in a CRISPR-CATCH reconstruction pipeline, available at https://github.com/siavashre/CRISPRCATCH.

1. To identify the regions of interest, we ran PrepareAA (https://github.com/jluebeck/PrepareAA) (version 0.931.4) and AmpliconArchitect (version 1.2_r2, available from https://github.com/jluebeck/AmpliconArchitect) on two public bulk SNU16 WGS datasets (SRX5466612^36^; SRR530826, Genome Research Foundation) and found comparable graphs in both. We used PrepareAA with BWA-MEM^26^ (version 0.7.12-r1039) to align reads to hg19 and CNVKit^37^ (version 0.9.7) to generate seed regions having copy number (CN) > 5. These regions were provided to AA, which constructed a CN-aware breakpoint graph. The genome regions AA included in the graph were converted to bed format and used as the seed regions in the analysis of each PFGE band, so that the regions studied were always consistent between bands.
2. Using WGS reads generated from CRISPR-CATCH-purified DNA, for each band we next aligned to the hg19 reference genome using PrepareAA which included BWA MEM and a PCR-duplicate removal step (using samtools^32^ version 1.3.1), and we also made estimates of insert size distribution using Picard (version 2.25.6) for quality control purposes.
3. The aligned PFGE data and seed regions identified from bulk sequencing were provided to AmpliconArchitect (version 1.2_r2) to construct the CN-aware breakpoint graph, using non-default arguments –downsample -1 –pair_support 2 –no_cstats – insert_sdevs 8.5. The –insert_sdevs parameter allows for larger insert size variation without forming breakpoints from read pairs marked as discordant, as we found high insert size variance occurred frequently in DNA extracted from the gels. Following AA, we ran a script on the resulting CN-aware breakpoint graph to filter non-foldback graph edges joining regions smaller than 1 kbp from the graph, representing potential unfiltered artifact edges arising from overdispersion in insert size variance, in order to reduce the complexity of the graph when performing pathfinding. Since the edges removed joined regions not more than 1 kbp apart and did not lead to changes in the orientation of the genome, this step had a negligible effect on the resulting paths. This utility for filtering AA graphs is made available as part of PrepareAA (graph_cleaner.py).
4. Central to the method for ecDNA reconstruction is the assumption that a single ecDNA is being analyzed within the graph, and as a result the estimated genomic copy numbers should closely relate to the number of times a segment appears within the ecDNA. We termed the number of times a segment appeared within a single ecDNA as the “multiplicity” of a genomic segment. The path finding method first removes low CN elements from the graph representing the background genome and contamination from incomplete separation of ecDNAs (i.e., remove segments with CN below 20% of the maximum CN of all segments having length > 100bp, or below 10% of the maximum, if the maximum CN is >10000). In the remaining segments, we assumed that the majority of segments appeared once within an ecDNA. We assumed that ecDNA where the majority of segments are present more than once would reflect cases where two or more ecDNA were present instead of one. Thus, to compute the multiplicity of each graph segment, the method computes the 40^th^ percentile of the remaining graph segment copy numbers and assigns that copy number, *S1,* to multiplicity = 1. For each segment, *i*, in the graph, we computed its multiplicity, *M(i)* as.

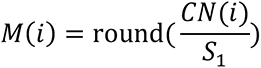
5. To find paths in the graph which represented candidate ecDNA structures, we used an exhaustive search constrained by the multiplicities of the segments and (if available) the estimated maximum molecule size suggested by the CRISPR-CATCH data. Candidate ecDNA structures are determined through a constrained depth-first search (DFS) approach, which attempts to identify paths in the graph, and performs the process starting at every segment in the graph assigned a non-zero multiplicity. During the search, the length of the path (in base pairs) must remain less than the maximum allowed length (*L*). For every segment *i*, appearing *ni* times in the path, *ni ≤ M(i)*. The DFS recursion terminates if either constraint is violated, and the current path is scored as ∑_“_ *n*_”_. The path is compared against the current best path (initiated as an empty path with score 0) and updated if it scores higher. Both the best-scoring cyclic paths as well as the best-scoring paths regardless of cyclic status are returned after removing all duplicate (identical) paths from the collection of best-scoring paths. This utility is also individually available from PrepareAA (plausible_paths.py).
6. We found a number of features of both the breakpoint graph and the reconstructions to be informative about the quality of the data in the band. We developed quality annotations reported along with each reconstruction to provide users with annotations about the confidence of the reconstruction. We note that CN-aware breakpoint graphs derived from NGS data may contain a number of error sources including missing edges between graph segments and incorrect estimation of copy numbers (leading sometimes to incorrect estimation of multiplicity). The method applies the following filters.

a) In the amplicon region analyzed by AA, the total amount of amplified material (non-zero multiplicity) should not significantly exceed the maximum estimated molecular size of the band (if provided). We used a cutoff such that amplicons with 1.4x the maximum estimated molecular size of the band were flagged for low quality (incomplete separation of ecDNA).
b) Changes in multiplicity must be accompanied by one or more breakpoint junctions, and thus for a breakpoint graph with |*e|* total edges, amplicons where

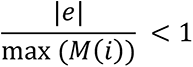
were flagged for low quality (missing graph edges).
c) We defined a root mean square residual for the unexplained copy numbers of *M(i)*. In a given path, for each segment *i*, having *ni* occurrences in the path, the root mean square residual was defined as

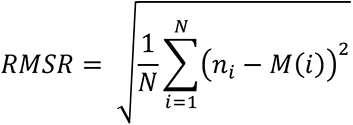
where *N* is the number of segments having non-zero multiplicity in the graph. We set a default cutoff such that amplicons with RMSR > 0.9 were flagged as low quality (too many amplified graph segments having incompletely used multiplicity).
d) To assess how tightly segment copy numbers could be segregated by segment multiplicity, we computed the Davies-Bouldin index^38^ (DBI) on the clusters of copy numbers. Each cluster was comprised of all segment copy numbers assigned to a multiplicity (singleton clusters excluded), and the centroid of the cluster was the mean CN for the cluster. Amplicons where the DBI was > 0.3 were flagged as low quality due to noisy copy number estimation.
e) If a minimum molecular size for the band was given, we flagged reconstructions which fell below that 90% of that value as low quality as they reflected incomplete reconstructions.
f) If no segment in the reconstruction overlapped the CRISPR-Cas9 target site, we flagged it as being low quality as it was either an incomplete reconstruction, or the incorrect amplicon was detected.
7. Since the reconstructed paths are reported in the textual AA_cycles.txt format, the method also provides automated circular visualizations of the structures and the WGS coverage tracks which are generated by CycleViz (https://github.com/jluebeck/CycleViz) (version 0.1.0).

### Optical Mapping

Optical maps from SNU16 cells were generated as follows: ultra-high molecular weight (UHMW) DNA was extracted from 1.5 million frozen cells preserved in DMSO following the manufacturer’s instructions (Bionano Genomics, #30398, 80042). Briefly, cells were digested with Proteinase K (Puregene #158920) and RNAse A (Puregene #158922) and then the DNA was precipitated with isopropanol and bound with nanobind magnetic disks. Bound UHMW DNA was resuspended in elution buffer (EB) and quantified with Qubit dsDNA BR assay kit (ThermoFisher Scientific, Q32850). The final DNA concentration was initially too high (280 ng/µl), therefore UHMW DNA was further diluted with EB, gently mixed with a wide-bore tip five times, and allowed to relax at room temperature for two days. Upon resting, UHMW DNA was diluted to 110 ng/µl in EB and DNA labeling was performed following the manufacturer’s instructions (Bionano Genomics, #30206, 80005). Standard Direct Labeling Enzyme 1 (DLE-1) reactions were carried out using 750 ng of purified UHMW DNA. Using the Qubit dsDNA HS assay kit (ThermoFisher Scientific, #32854), the final labeled DNA concentration was determined as 5.4 ng/µl with a coefficient of variation of 0.026. The fluorescently labeled DNA molecules were loaded onto the Saphyr Chip G2.3 (Bionano Genomics, #20366, 30142) and were imaged sequentially across nanochannels on a Saphyr instrument (Bionano Genomics, #90023). An effective genome coverage of approximately 340X, using molecules >= 150 kbp (molecule N50 of 0.2505 Mbp) was achieved.

*De novo* assembly of SNU16 was performed with Bionano’s *de novo* assembly pipeline (Bionano Solve v3.6, #90023) using standard haplotype aware arguments. With the Overlap-Layout-Consensus paradigm, pairwise comparison of DNA molecules of approximately 130X coverage was used to create a layout overlap graph, which was then used to generate the initial consensus genome maps, which had a contig N50 of 50 Mbp. By realigning molecules to the genome maps (P value cutoff of <10^-12^), and by using only the best matched molecules, a refinement step was done to refine the label positions on the genome maps and to remove chimeric joins. Next, during an extension step, the software aligned molecules to genome maps (P<10^-12^), and extended the maps based on the molecules aligning past the map ends. Overlapping genome maps were then merged (P<10^-16^). These extension and merge steps were repeated five times before a final refinement (P<10^-12^) was applied to “finish” all genome maps.

### Validating candidate structures with optical mapping

To validate candidate ecDNA paths we used long-range optical mapping (OM) data. Previously, we developed a method, AmpliconReconstructor (AR)^7^, which uses OM data and AA’s outputs as inputs. AR attempts to identify paths within the breakpoint graph supported by OM contigs. In the CRISPR-CATCH data, the graphs may have many smaller segments than the graphs derived from bulk WGS, due to noisier CN profiles in the gel-extracted DNA, or due to highly complex ecDNA structures having very dense breakpoints. In these cases, large numbers of graph segments may be too small to reliably align against OM contigs using AR’s standard approach. Thus, we added a method to AR whereby the user can provide an AA-formatted cycles.txt file containing the full candidate path. The individual segments are combined into a single long genome sequence, then converted to an *in silico* digested OM sequence. The combined candidate OM sequence is then aligned with OM contigs using AR’s SegAligner method. The resulting candidate sequence alignment label indices are mapped back to the original uncombined candidate path. Contig alignments for the candidate path were combined if necessary and visualized using CycleViz. The candidate path alignment utility added to AR for this analysis is available at https://github.com/jluebeck/AmpliconReconstructor.

## Data Availability

Sequencing data generated in this study are deposited in SRA under BioProject accession PRJNA777710.

## Code Availability

Custom code to perform reconstructions of candidate ecDNA structures from CRISPR-CATCH data is available at https://github.com/siavashre/CRISPRCATCH.

## Acknowledgements

We thank members of the Chang and Bafna laboratories for discussions. H.Y.C. was supported by NIH R35-CA209919 and RM1-HG007735. K.L.H. was supported by a Stanford Graduate Fellowship. H.Y.C. is an Investigator of the Howard Hughes Medical Institute.

## Author Contributions

K.L.H. and H.Y.C. conceived the project. K.L.H. performed experiments for CRISPR-CATCH method development for ecDNA purification and analyses of genetic variants and epigenetic features from short-read sequencing and nanopore sequencing data. J.L. and S.R.D. analyzed short-read sequencing and optical mapping data for amplicon reconstruction. C.C. and J.A.L. generated optical mapping data and provided *de novo* assembly and rare variant analysis results. W.J.G. advised on single molecule sequencing. P.M., V.B. and H.Y.C. guided data analysis and provided feedback on experimental design. K.L.H. and H.Y.C. wrote the manuscript with input from all authors.

## Materials & Correspondence

Correspondence and requests for materials should be addressed to Howard Y. Chang.

---

**Extended Data Figure 1.**
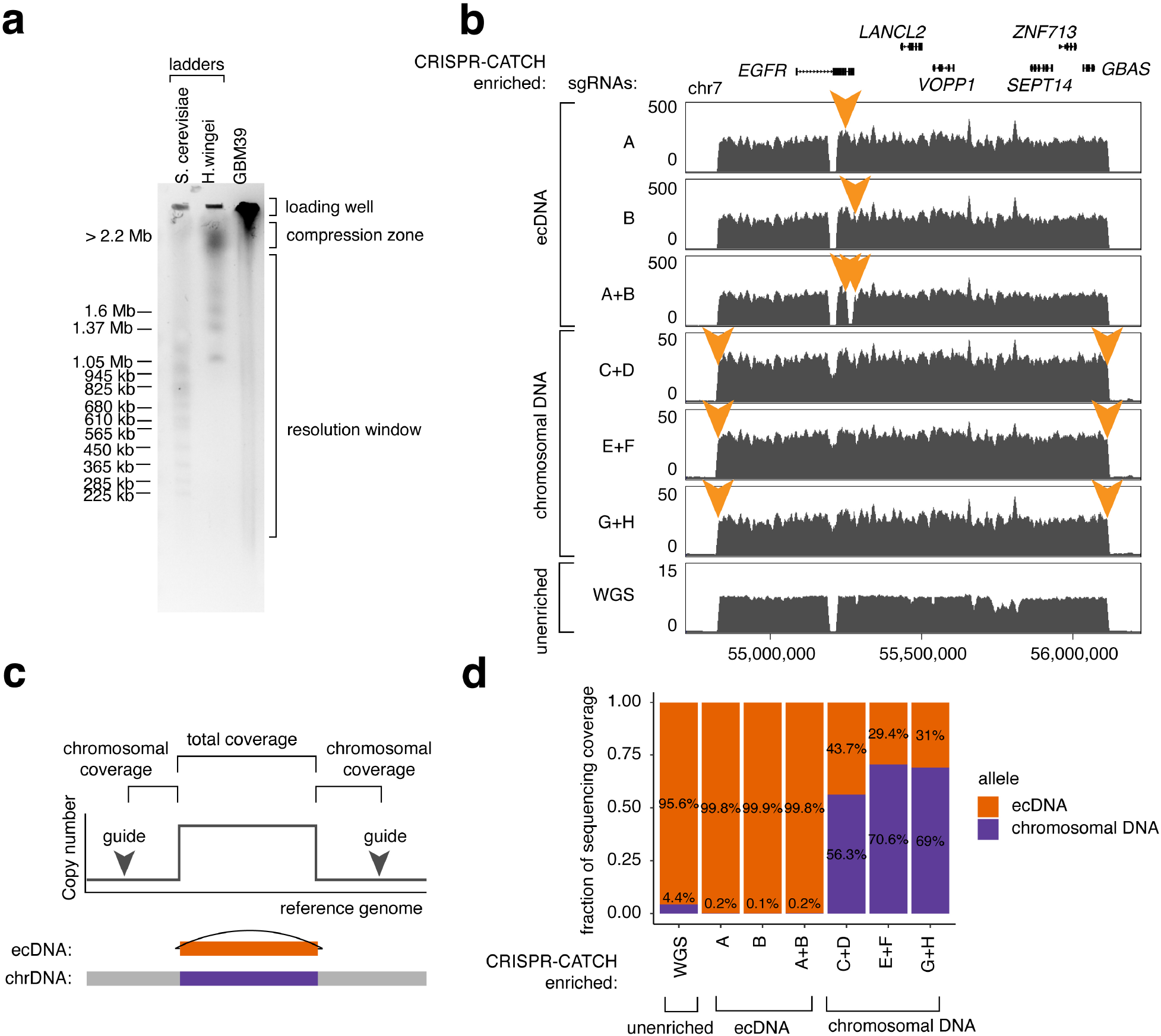
Enrichment of circular ecDNA by CRISPR-CATCH. **(a)** A representative PFGE image showing size ladders and GBM39 ultra-high molecular weight (UHMW) genomic DNA without *in-vitro* CRISPR-Cas9 linearization. UHMW DNA was trapped in the loading well and the upper compression zone. **(b)** Full sequencing tracks showing coverage for purified ecDNA and its chromosomal locus at the EGFR amplified region compared to WGS. Zoomed-in tracks are shown in Figure 1f. Orange arrows indicate locations of sgRNA targets. **(c)** Chromosomal overhangs from chromosome-targeting guides (guides C-H) outside of the ecDNA-amplified region were used for calculating sequencing coverage of the chromosomal allele. The mean coverage of the 5’ and 3’ chromosomal overhangs was calculated. The coverage of ecDNA alleles was calculated by subtracting chromosomal coverage from total coverage in the ecDNA-amplified region. **(d)** Relative sequencing coverage of chromosomal DNA and ecDNA alleles in WGS or CRISPR-CATCH samples.

**Extended Data Figure 2.**
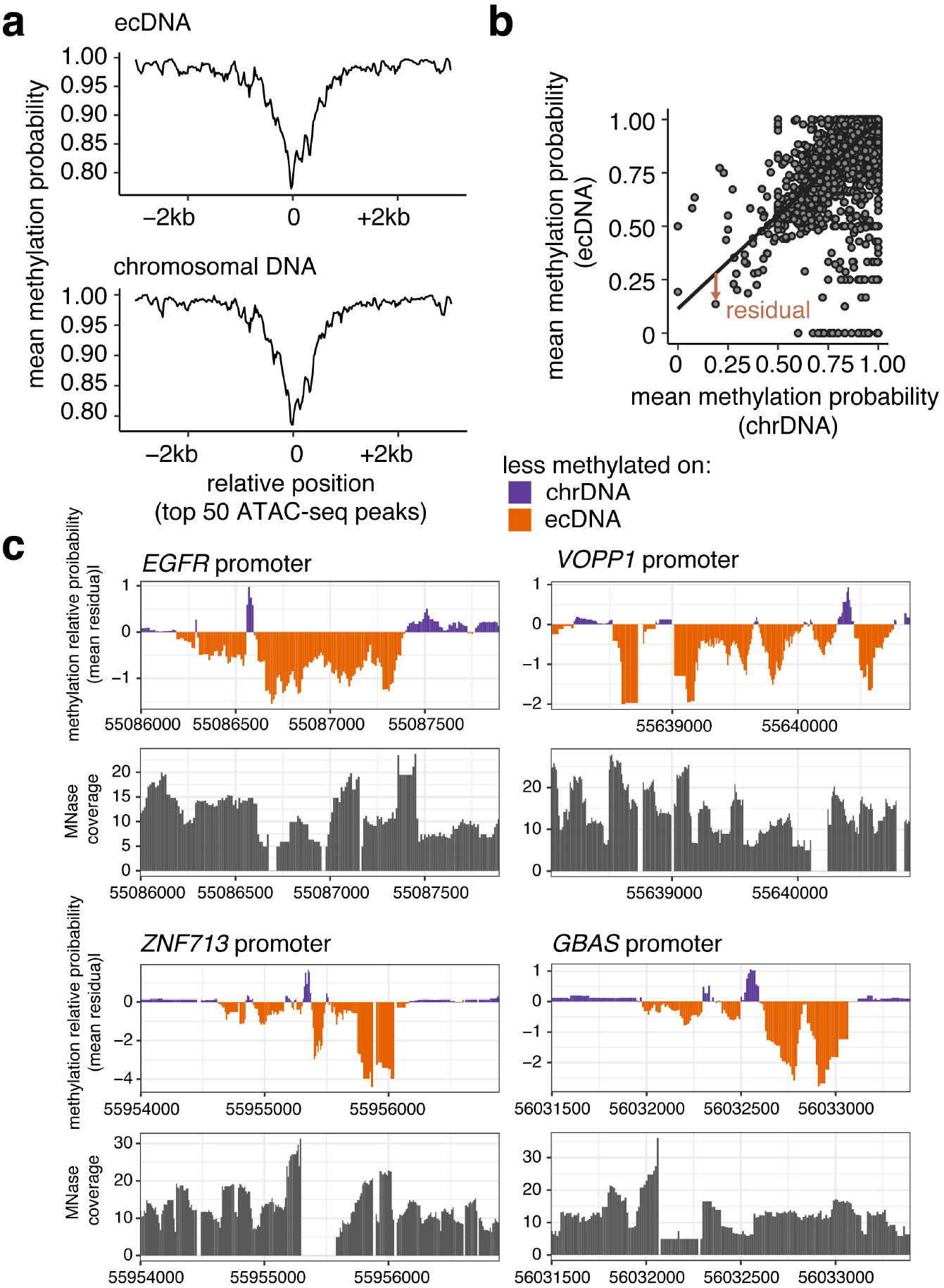
Quantification of 5mC-CpG methylation probability of ecDNA and the native chromosomal locus. **(a)** Aggregated CpG methylation probability of ecDNA and chromosomal DNA at the top 50 ATAC-seq peaks with highest coverage in the amplified region. Mean methylation frequencies were calculated in 100- bp windows sliding every 10 bp. **(b)** Linear regression model of mean methylation probabilities of ecDNA vs chromosomal DNA. Mean methylation probabilities were calculated in 100-bp windows sliding every 10 bp in the ecDNA-amplified region. Each point represents a window mean. Brown arrow demonstrates the standardized residual of a data point from the regression line. **(c)** Relative CpG methylation of ecDNA compared to the chromosomal locus and nucleosome positioning by MNase-seq, zooming into indicated gene promoters. Regions shown correspond to differentially methylated regions in **Figure 3d,e**. Mean methylation frequencies and MNase-seq coverage were calculated in 100-bp windows sliding every 10 bp. Relative frequencies were quantified from standardized residuals for a linear regression model for mean frequencies on ecDNA vs chromosomal DNA (Methods).

**Extended Data Figure 3.**
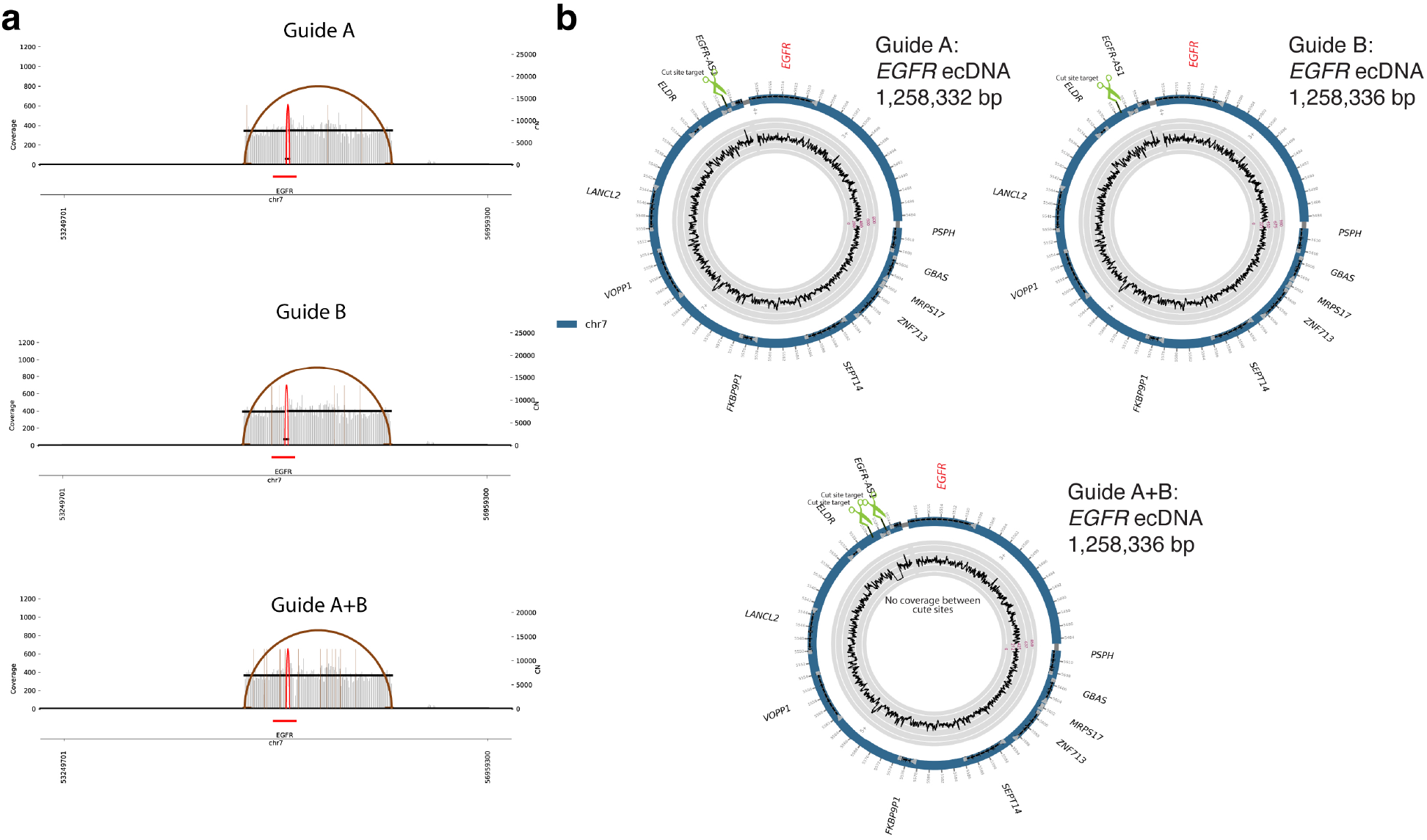
Reconstruction of a 1.258 Mb ecDNA from GBM39 neurospheres. **(a)** AmpliconArchitect breakpoint graphs for CRISPR-CATCH-purified ecDNAs using guides A and/or B as in Figure 1 (guide sequences in **Supplementary Table 1**). **(b)** Reconstructed ecDNA circles from CRISPR-CATCH data using independent sgRNAs showing equivalent ecDNA structures (outer rings; thin grey bands mark connections between sequence segments). Sequencing coverage is shown along the reconstructed circle (inner rings). Green scissors mark sgRNA target sites. Coordinate tick marks are printed in 10-kb units. AmpliconArchitect segment IDs and orientations are annotated.

**Extended Data Figure 4.**
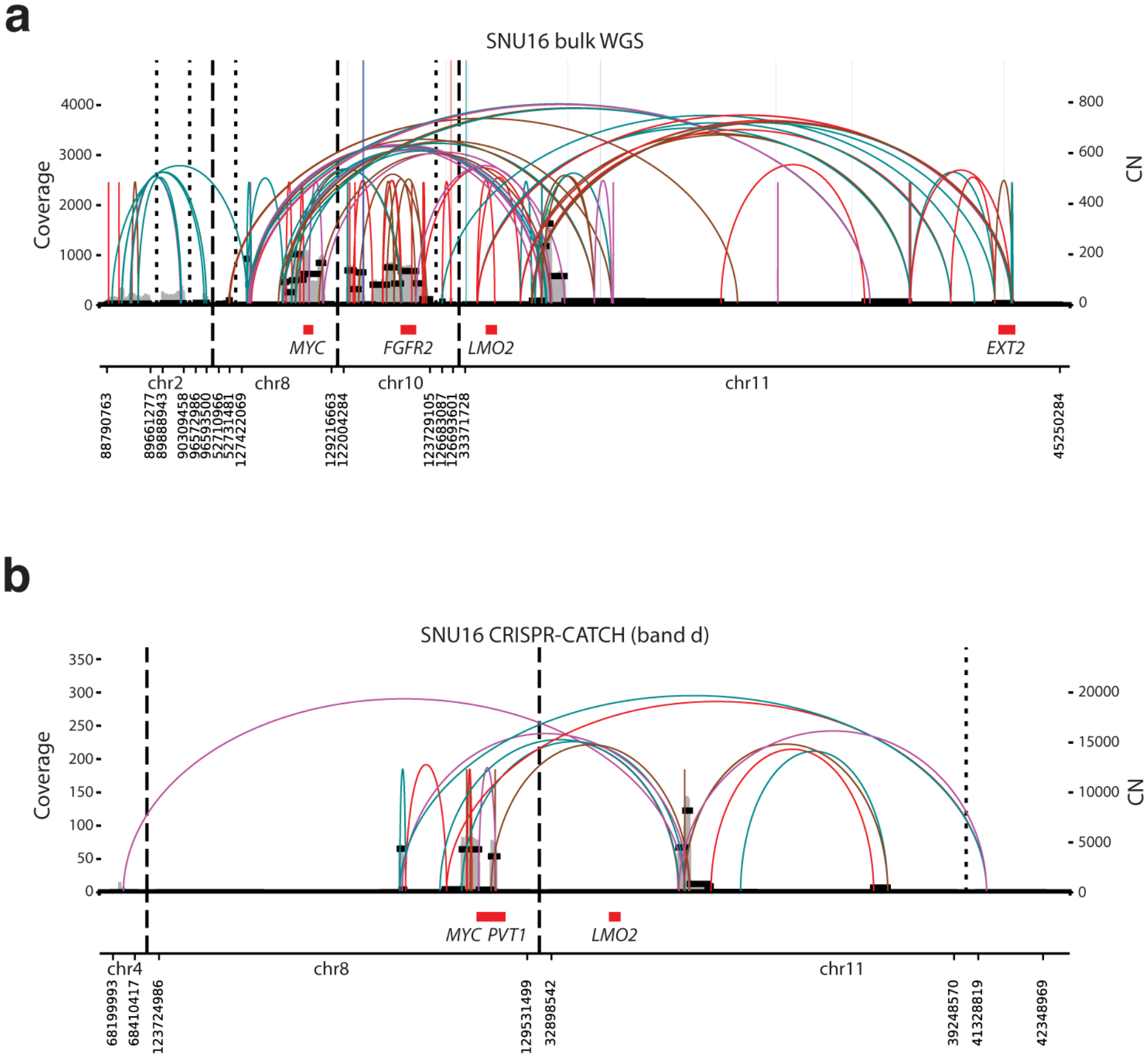
CRISPR-CATCH enables disambiguation of heterogeneous sequence rearrangements on individual ecDNA species. **(a)** AmpliconArchitect breakpoint graph from bulk WGS of stomach cancer SNU16 cells showing significantly amplified sequences from chromosomes 8, 10, and 11. **(b**) An example of an AmpliconArchitect breakpoint graph for a CRISPR-CATCH-separated ecDNA species (band “d”) from SNU16 cells showing greatly simplified breakpoints connecting only sequences from chromosomes 8 and 11. Gray vertical lines represent genomic coverage from WGS data and black horizontal lines indicate the estimated copy number of the region. Colored arcs represent breakpoint junctions, and the orientation of those junctions is specified by the color. Red and brown arcs preserve the orientation of the genome, with red reflecting breakpoints supported by reads in the proper orientation and brown reflecting breakpoints supported by reads in the everted orientation. Teal and magenta arcs indicate breakpoints leading to a change in genome orientation before and after the breakpoint where teal breakpoints are supported by both paired-end reads mapping to the forward strand and magenta breakpoints are supported by both paired-end reads mapping to the reverse strand.

**Extended Data Figure 5.**
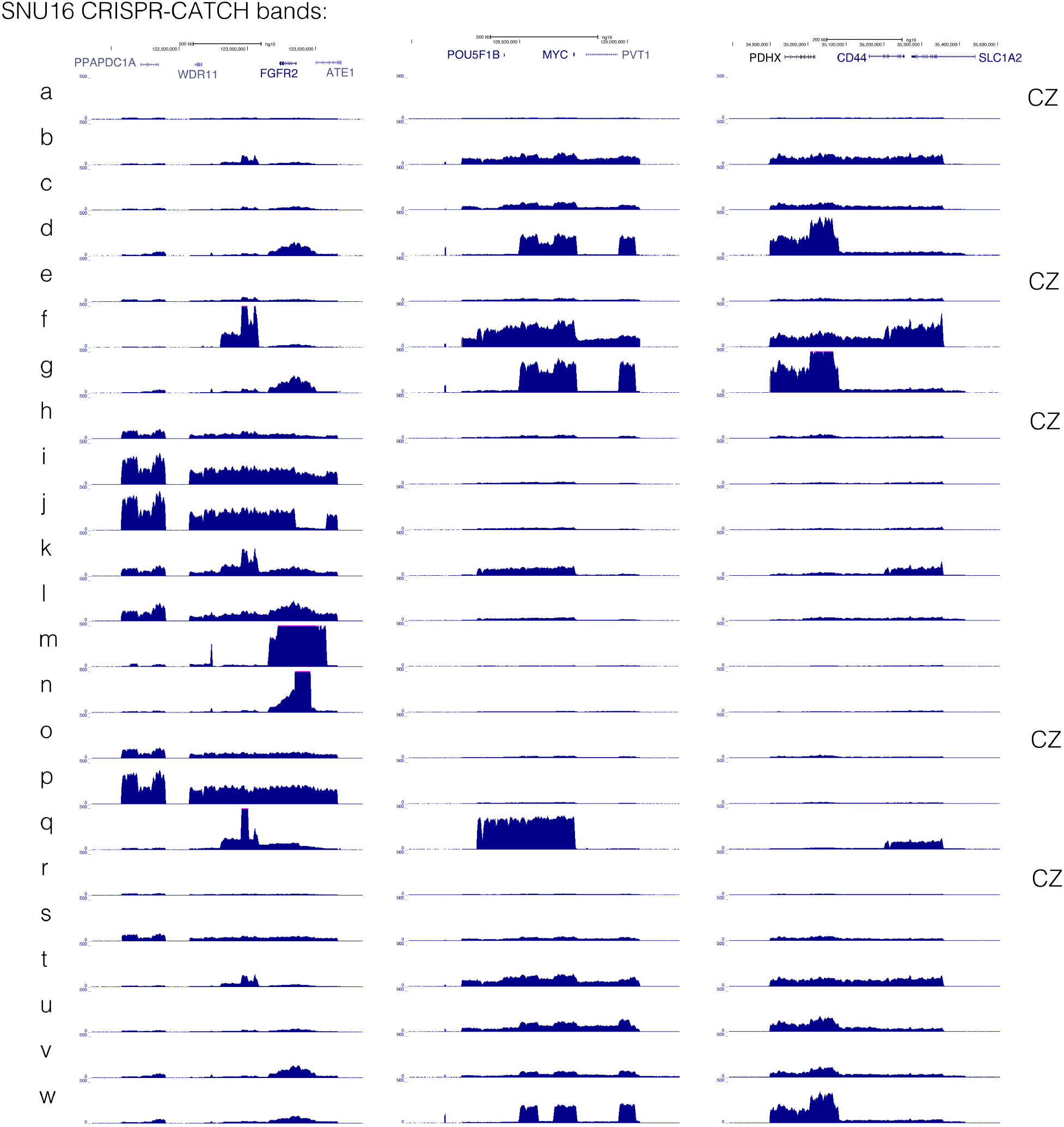
Enrichment of multiple ecDNA species from the SNU16 stomach cancer cell line. Short-read sequencing coverage tracks of multiple ecDNA species from SNU16 cells after CRISPR-CATCH purification at the *FGFR2*, *MYC* and *CD44* loci. Bands a-w correspond to extracted bands shown in Figure 4b. Bands corresponding to unresolved DNA content in the compression zone are labeled CZ.

