## Supplementary Information for "Targeted profiling of human extrachromosomal DNA by CRISPR-CATCH"

**Table of Contents**

**Supplementary Figure 1.** Raw PFGE images corresponding to Figures 1c, 3, 4b, and Extended Data Figure 1a.

**Supplementary Table 1:** sgRNA sequences.

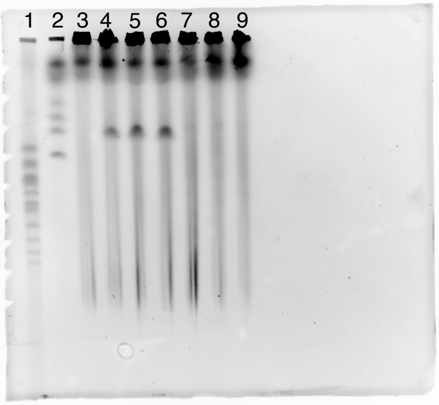


1. CHEF DNA Size Marker, 0.2–2.2 Mb, S. cerevisiae Ladder
2. CHEF DNA Size Marker, 1–3.1 Mb, H. wingei Ladder
3. no treatment
4. guide A
5. guide B
6. guide A+B
7. guide C+D
8. guide E+F
9. guide G+H

Raw image of PFGE agarose gel for CRISPR-CATCH for GBM39 cells. Image was cropped to remove extra white space and ladders, contrast was adjusted to make bands more visible. Corresponds to **Figure 1c**.

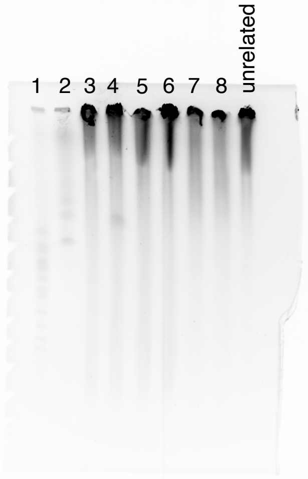


1. CHEF DNA Size Marker, 0.2–2.2 Mb, S. cerevisiae Ladder
2. CHEF DNA Size Marker, 1–3.1 Mb, H. wingei Ladder
3. GBM39 no treatment
4. GBM39 guide A
5. GBM39 guide E
6. GBM39 guide E+F
7. Jurkat no treatment
8. Jurkat guide A

Raw image of PFGE agarose gel for CRISPR-CATCH for indicated cell lines. Image was cropped to remove extra white space and ladders, contrast was adjusted to make bands more visible. Lanes 4/6 correspond to purified ecDNA and chromosomal DNA used in **Figure 3**, respectively.
Lanes 7/8 correspond to **Figure 1c**.

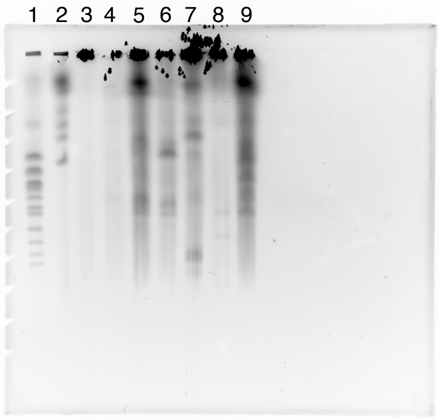


1. CHEF DNA Size Marker, 0.2–2.2 Mb, S. cerevisiae Ladder
2. CHEF DNA Size Marker, 1–3.1 Mb, H. wingei Ladder
3. no treatment
4. guide 7
5. guide 3
6. guide 5
7. guide 17
8. guide 18
9. guide 82

Raw image of PFGE agarose gel for CRISPR-CATCH for SNU16 cells. Image was cropped to remove extra white space and ladders, contrast was adjusted to make bands more visible. Corresponds to **Figure 4b**.

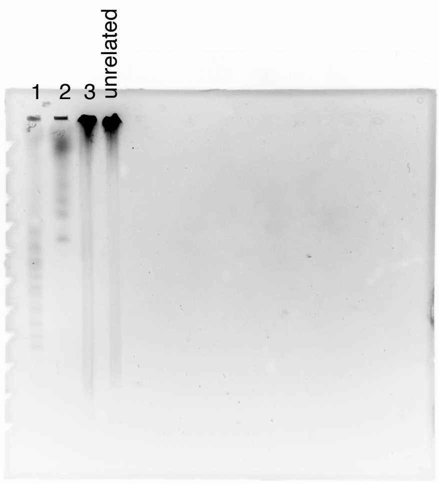


1. CHEF DNA Size Marker, 0.2–2.2 Mb, S. cerevisiae Ladder
2. CHEF DNA Size Marker, 1–3.1 Mb, H. wingei Ladder
3. no treatment

Raw image of PFGE agarose gel for untreated GBM39 cells. Image was cropped to remove extra white space. Corresponds to **Extended Data Figure 1a.**

### Supplementary Tables

#### Supplementary Table 1.

| ID | gRNA_sequence | gRNA_info |
|---|---|---|
| 1 | TGGCGCAGTTATGCTTTAAC | EGFR guide A, used in GBM39 experiments |
| 2 | GGATCTACTTGGCACTCGCT | EGFR guide B, used in GBM39 experiments |
| 3 | CAATACCGCACTCAATGTCA | EGFR guide C, used in GBM39 experiments |
| 4 | ACAAACCGCGAGATCAGGGG | EGFR guide D, used in GBM39 experiments |
| 5 | ACGTTAAAAAGCTGTCGCGC | EGFR guide E, used in GBM39 experiments |
| 6 | TCCCGTGCGCGATGACGACA | EGFR guide F, used in GBM39 experiments |
| 7 | TAAACCACGGAAGCGGCGGC | EGFR guide G, used in GBM39 experiments |
| 8 | GCCTTGTCGTCATCGCGCAC | EGFR guide H, used in GBM39 experiments |
| 9 | CCAGCAATCGTTAACCACTG | MYC guide 3, used in SNU16 experiments |
| 10 | CTTCGGGGAGACAACGACGG | MYC guide 5, used in SNU16 experiments |
| 11 | GTGATATTTGAACCGCCCTG | MYC guide 7, used in SNU16 experiments |
| 12 | GGGGATTGGTACCGTAACCA | FGFR2 guide 17, used in SNU16 experiments |
| 13 | GAGGCGATAATATCAACATG | FGFR2 guide 18, used in SNU16 experiments |
| 14 | ATCATGTAGTATCCCCCACC | MYC guide 82, used in SNU16 experiments |
